# Gut microbiome features associated with *Clostridium difficile* colonization in puppies

**DOI:** 10.1101/599647

**Authors:** Alexander S. Berry, Denise Barnhart, Brendan J. Kelly, Donna J. Kelly, Daniel P. Beiting, Robert N. Baldassano, Laurel E. Redding

## Abstract

In people, colonization with *Clostridium difficile*, the leading cause of antibiotic-associated diarrhea, has been shown to be associated with distinct gut microbial features, including reduced bacterial community diversity and depletion of key taxa. In dogs, the gut microbiome features that define *C. difficile* colonization are less well understood. We sought to define the gut microbiome features associated with *C. difficile* colonization in puppies, a population where the prevalence of *C. difficile* has been shown to be elevated, and to define the effect of puppy age and litter upon these features and *C. difficile* risk. We collected fecal samples from weaned (n=27) and unweaned (n=74) puppies from 13 litters and analyzed the effects of colonization status, age and litter on microbial diversity using linear mixed effects models.

Colonization with *C. difficile* was significantly associated with younger age, and colonized puppies had significantly decreased bacterial community diversity and differentially abundant taxa compared to non-colonized puppies, even when adjusting for age. *C. difficile* colonization remained associated with decreased bacterial community diversity, but the association did not reach statistical significance in a mixed effects model incorporating litter as a random effect.

Even though litter explained a greater proportion (67%) of the variability in microbial diversity than colonization status, we nevertheless observed heterogeneity in gut microbial community diversity and colonization status within more than half of the litters, suggesting that the gut microbiome contributes to colonization resistance against *C. difficile*. The colonization of puppies with *C. difficile* has important implications for the potential zoonotic transfer of this organism to people. The identified associations point to mechanisms by which *C. difficile* colonization may be reduced.

## Introduction

*Clostridium difficile* is a spore-forming anaerobic, gram-positive bacillus that is the leading cause of antibiotic-associated and nosocomial diarrhea in humans and a significant enteric pathogen in many species of animals. Administration of antibiotics is the primary risk factor for the development of *C. difficile* infection (CDI). However, patients can develop CDI outside of a healthcare facility without the prior use of antibiotics, and community-acquired CDIs are now thought to account for one quarter of infections (1, 2).

The source of community-acquired infections has not been definitively established. People asymptomatically colonized with *C. difficile* are potential reservoirs (3), but zoonotic, environmental, and food-borne transmission to people has also been posited. The presence of *C. difficile* in companion animals has been documented since the 1980’s, and dogs and cats were posited as a potential reservoir species as early as 1983 (4). Given the close contact between people and their pets, colonized or infected companion animals may represent an important transmission source for this pathogen. As in other species of animals (5–8), including human infants (9–11), *C. difficile* is highly prevalent in the feces of puppies (12–14). Understanding how colonization is regulated in puppies might reduce their colonization with *C. difficile* and the potential transmission to pet owners.

The role of the commensal gut microbiota in *C. difficile* colonization resistance has been demonstrated in people (15–19) and in certain species of animals (20–22). Human subject and animal model studies suggest key microbiome features, including community diversity and specific taxa, are involved in protection against *C. difficile*. No such association has been demonstrated in dogs, and studies of the association between the administration of antibiotics (and the consequent disruption of the gut microbiota) and *C. difficile* colonization/infection in dogs have yielded mixed results (23, 24). The evolution of the neonatal canine gut microbiome has been described, with increasing diversity and taxonomic shifts occurring with increasing age (25). As has been found in human infants (17), it is possible that certain taxonomic patterns and a lack of microbial community diversity in the gut may be associated with a lack of colonization resistance to *C. difficile*.

The objective of this study was to define the gut microbiome features associated with *C. difficile* colonization in puppies and to define the effects of puppy age and litter on the risk of colonization. The results could contribute to a better understanding of *C. difficile* colonization in puppies and their potential to serve as a reservoir for this pathogen.

## Materials and Methods

### Samples

Fecal samples were obtained by pet owners bringing their puppies to the pediatric service at the Veterinary Hospital of the University of Pennsylvania and by breeders in the greater Philadelphia area who collected fecal samples from their puppies and shipped them on ice overnight to the laboratory. All puppies were healthy at the time of sampling, and none had received antimicrobial therapy. After collection, samples were split into sterile cryogenic vials. One aliquot was processed for culture within 24 hours, while others were stored at −80°C and processed subsequently in batch for the 16S ribosomal RNA (rRNA) sequencing.

Frozen samples were thawed only once prior to processing. This study was approved by the Institutional Animal Care and Use Committee of the University of Pennsylvania (protocol number 806539).

### Anaerobic culture and toxigenic testing

A 0.5 g pellet of formed fecal sample was mixed with 0.5 ml of 100% ethanol. The mixture remained for 60 minutes at room temperature before being inoculated on BBL™ CDSA*/Clostridium difficile* selective agar (BD; Sparks, Maryland, USA) and Columbia CNA agar (Remel; Lenexa, KS, USA). Inoculated plates were incubated at 35°C under anaerobic growth conditions for seven days and checked for growth every other day. Suspect colonies were identified and isolated. Isolates were confirmed to be *C. difficile* by Maldi-TOF MS identification and/or RapID™ ANA II System (ThermoFisher Scientific, USA). Confirmed isolates of *C. difficile* were inoculated into BHI broth and/or cooked meat broth to induce toxin production. The broth was incubated anaerobically at 35°C for 48 hours. The supernatant was collected and tested by EIA (TechLab *C. difficile* Tox A/B II™) for toxin production.

### 16 S sequencing

DNA was extracted from the fecal samples using Qiagen Power Soil DNA Extraction Kit (Qiagen, Hilden, Germany) using 0.25 g of each fecal pellet as input. Extraction and PCR blanks were used to control for environmental contamination and mock communities were used to control for contamination across wells. The V4 region of the 16S rRNA gene was amplified using barcoded primers for use on the Illumina platform (26). The concentration of each PCR product was determined using a PicoGreen assay, and samples were normalized to equal amounts and pooled. Sequencing was performed using 250-base paired-end chemistry on an Illumina MiSeq instrument with an average read depth of 49,436 reads per sample. Three samples were dropped due to low read depth (<4000 reads per sample), raising the average read depth to 50,860 reads per sample. Sequences were demultiplexed using the Quantitative Insights into Microbial Ecology (QIIME2) software (27), and denoised using DADA2 (28). Sequences were aligned using Maaft (29) and phylogenetic reconstruction was performed using Fasttree (30). Finally, sequences were rarefied to 11,700 reads per sample for calculating alpha- and beta-diversity metrics.

### Analysis

The effects of age and litter on culture status were analyzed by logistic regression. Metrics of alpha and beta diversity of the fecal microbiome were calculated using the qiime diversity core-metrics-phylogenetic function in qiime2 and visualized using QIIME2 and Emperor (31).

The alpha diversity was calculated for each sample using the Shannon index. Differences in alpha diversity between *C. difficile*-infected and uninfected puppies were assessed using (1) univariable linear regression (N=98), (2) linear regression controlling for puppy age (N=98), and (3) a linear mixed effects model (LMM) on all unweaned puppies, controlling for age and using litter as a random effect (N=70) using the lme4 package in R (32). The effect of *C. difficile* colonization status on microbiome alpha diversity was assessed by comparing the likelihoods of the LMM with and without the fixed effect of *C. difficile* infection status using an analysis of variance. Finally, the effect of *C. difficile* toxigenicity on alpha diversity among *C. difficile*-positive puppies was assessed using univariable linear regression (N=35).

The effect of *C. difficile* culture status on the per-specimen bacterial community diversity of the fecal microbiome was first assessed by univariable analysis. Univariable analysis was also performed to identify clustering of specimens by colonization status, using the PERMANOVA test applied to pairwise distances as determined by the beta diversity metrics Bray-Curtis, unweighted unifrac, and weighted unifrac. The effect of *C. difficile* culture status on beta diversity of the microbiome adjusted for puppy age and litter was assessed using mixed effects PERMANOVA. Age and culture status were considered fixed effects, while litter was considered a random effect. All comparisons were two-tailed, and P < 0.05 was considered to represent statistical significance. PERMANOVA tests were performed using the vegan package (33) as implemented in R v.3.5.2 (R Core Team, 2018). Principal coordinates analysis (PCoA) was performed using phyloseq (34) to visualize the clustering of samples by various parameters (*C. difficile* status, age, litter).

A taxonomic classifier trained on the GreenGenes database with 99% Operational taxonomic units (OTUs) was used to assign relative abundances of OTUs for each sample calculated at the genus level. The relative contributions of different microbial taxa that characterize the differences between *C. difficile* culture positive and negative puppies were assessed through linear discriminant analysis effect size (LEfSe) using the tools found at http://huttenhower.sph.harvard.edu/galaxy/. OTUs were filtered such that only those with >5% relative abundance in one or more samples and with LDA scores > 2.0 were considered to be significant. All plots were generated using the ggplot2 package in R (35).

## Results

### Subject characteristics and *C. difficile* status

A total of 101 samples were collected from puppies ranging in age from 2-28 weeks. Seventy-four of the samples were obtained from 13 different litters of puppies that were still with their dam, and 27 samples were obtained from older weaned puppies that had been placed with families. The distribution of age was bi-modal, with the age of unweaned puppies in litters being significantly lower (p=0.01) than that of the weaned puppies (Figure 1). The mean (SD) age of the unweaned puppies was 3.7 (0.8) weeks, whereas that of the weaned puppies was 11.4 (2.9) weeks. Litters ranged in size from 3 to 12 puppies, with a mean (SD) of 5.8 (2.9) puppies.

**Figure 1:**
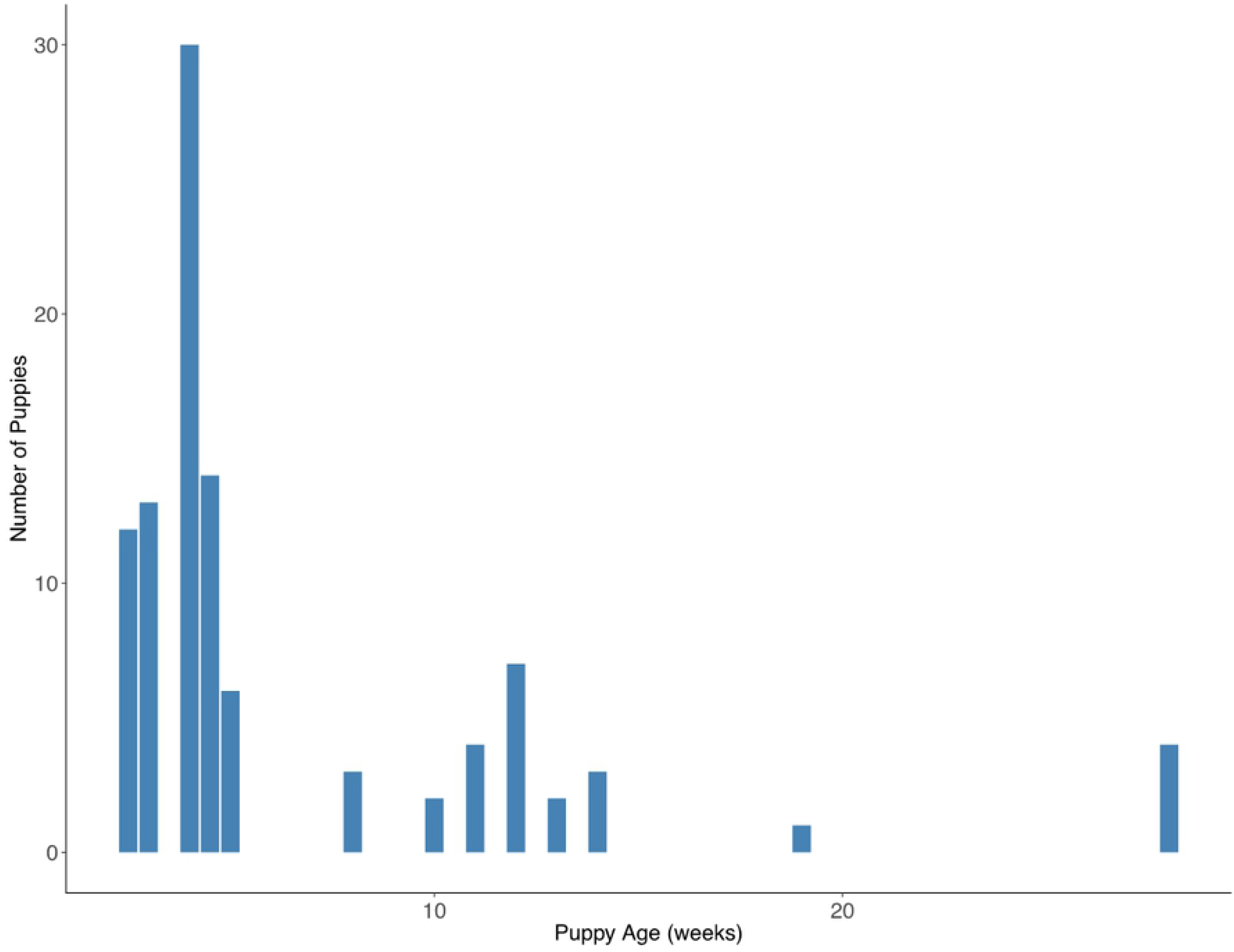
Distribution of the ages of puppies sampled in the greater Philadelphia region

Thirty-seven samples (36.3%) were culture-positive for *C. difficile*, and 19 (51%) of these *C. difficile* isolates were toxigenic. All of samples from the weaned puppies (n=27) were culture-negative. In 6 of the 13 litters of puppies, colonization status was the same for all puppies (i.e., all puppies within the litter were culture-negative or culture-positive). Age was significantly associated with culture status, with younger puppies being significantly more likely to harbor *C. difficile* (OR=0.46, p=0.004, 95% CI=0.27-0.78).

### Association between *C. difficile* status, age and microbiome diversity in all puppies

Complete 16 S sequencing was performed on 101 fecal samples. Three culture-negative samples were dropped from subsequent analyses because of low coverage. Alpha diversity was significantly lower (p<0.001) in the *C. difficile*-positive fecal samples than in the *C. difficile*-negative fecal samples (Figure 2). When adjusting for age, the effect of *C. difficile* status on microbial community diversity was mitigated but persistent (p-value increased 2 orders of magnitude from 1.6 e^−7^ to 6e^−5^). There was no difference in diversity between puppies colonized with toxigenic *C. difficile* and non-toxigenic *C. difficile* (p=0.66).

**Figure 2.**
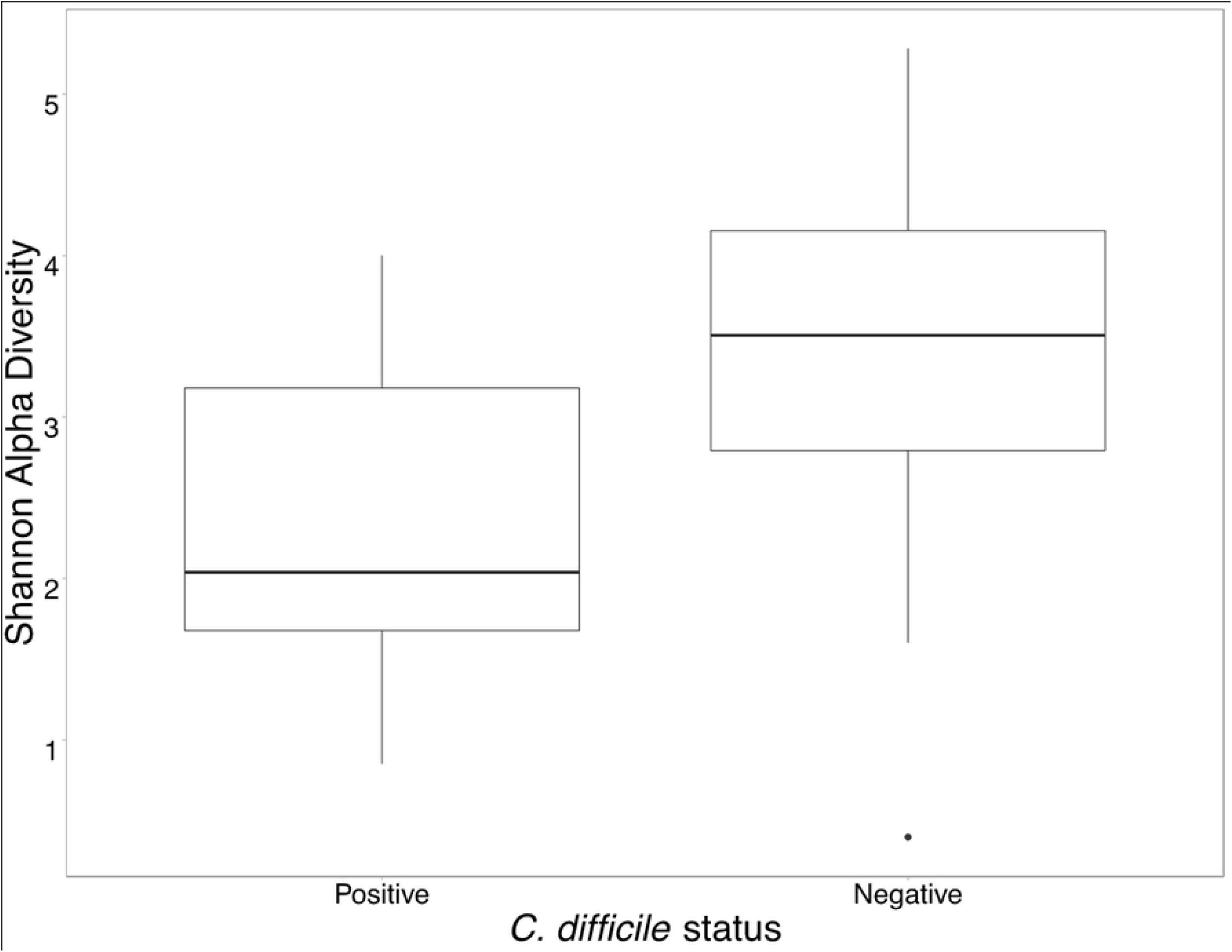
Boxplot of the Shannon diversity indices among *C. difficile*-positive puppies (left) and *C. difficile*-negative puppies (right). Boxes display the median, first and third quartiles, and whiskers extend to the minimum and maximum, while points represent outliers.

Beta diversity, or the dissimilarity between microbiome communities, was assessed using Bray-Curtis, weighted unifrac, and unweighted unifrac. Univariable analysis showed a significant difference between microbial communities using all three metrics (p=0.0001) even when controlling for age (p<0.0002) (Figure 3). The Bray-Curtis dissimilarity is summarized in a PCoA plot (Figure 3).

**Figure 3.**
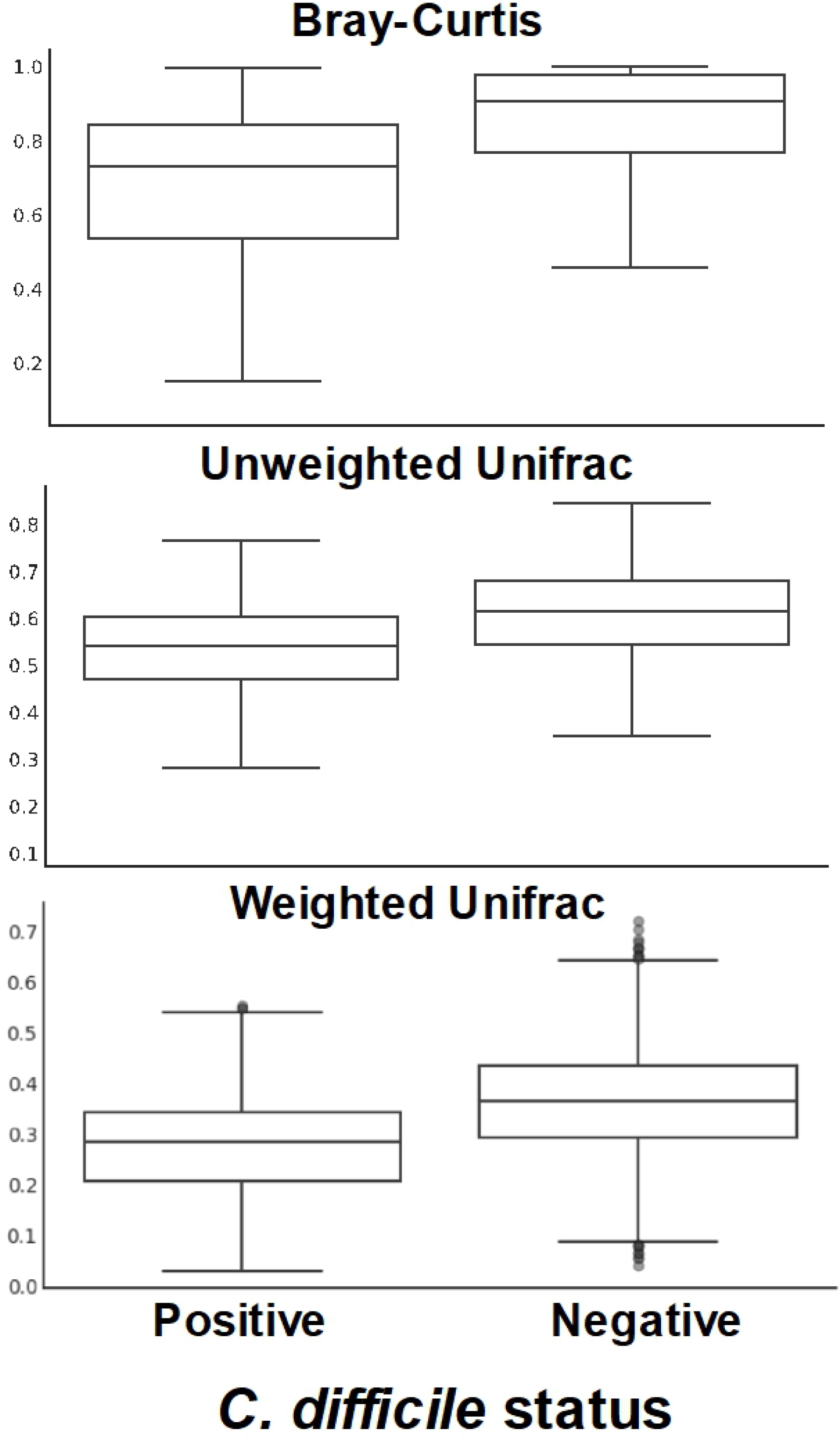
Box plots showing dissimilarity in bacterial communities in *C. difficile*-positive and *C. difficile*-negative fecal samples from puppies in the greater Philadelphia area. The dissimilarity among *C. difficile* positive puppies is displayed in the left boxplots and the dissimilarity between culture positive and negative puppies is displayed in the right boxplots. Boxes display the median, first and third quartiles, and whiskers extend to the minimum and maximum, while points represent outliers.

We found several taxa of bacteria to be differentially enriched in the *C. difficile*-positive and -negative samples. *C. difficile*-positive samples were enriched with members the *Escherichia, Bacteroides, Enterococcus* and *Parabacteroides* genera (Figure 4). Taxa from the *Escherichia* genus were found at relative abundance levels exceeding 10% in 48 samples and 50% in 15 samples. The relative abundance of *Escherichia* was associated with much of the clustering along the axis of principal component 1 (Figure 5). In contrast, *C. difficile*-negative samples were enriched with members of the *Prevotella, Megamonas*, and *Streptococcus* genera. Unweaned puppies that were not colonized with *C. difficile* had higher relative abundance of taxa from the *Clostridia* genera than unweaned puppies that were colonized with *C. difficile*.

**Figure 4.**
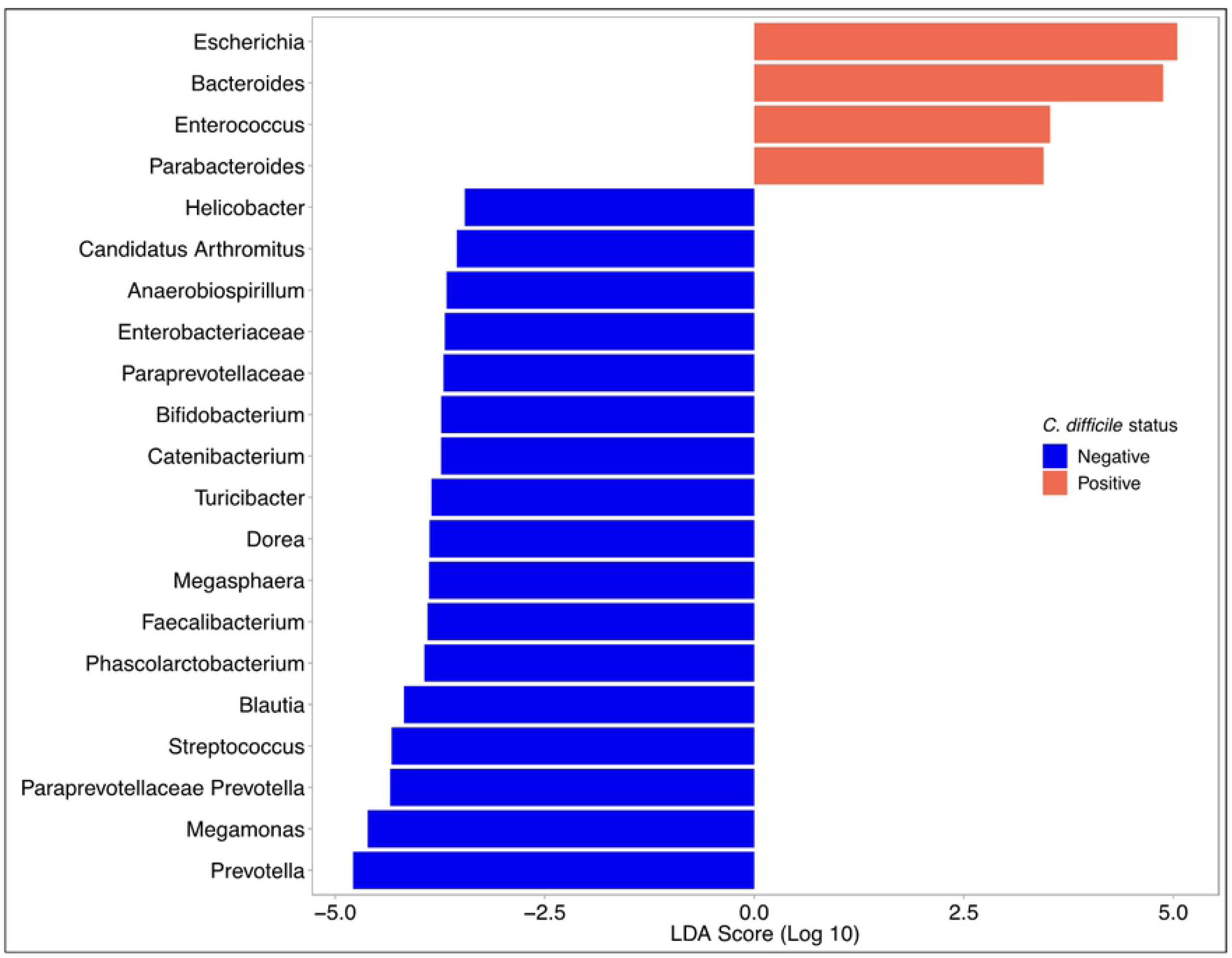
Linear discriminant analysis effect size analysis shows genera of bacteria that are differentially expressed in the *C. difficile*-positive and *C. difficile*-negative fecal samples from puppies in the greater Philadelphia area. Only organization taxonomic units with >5% relative abundance in one or more samples and with LDA scores > 2.0 are shown.

**Figure 5.**
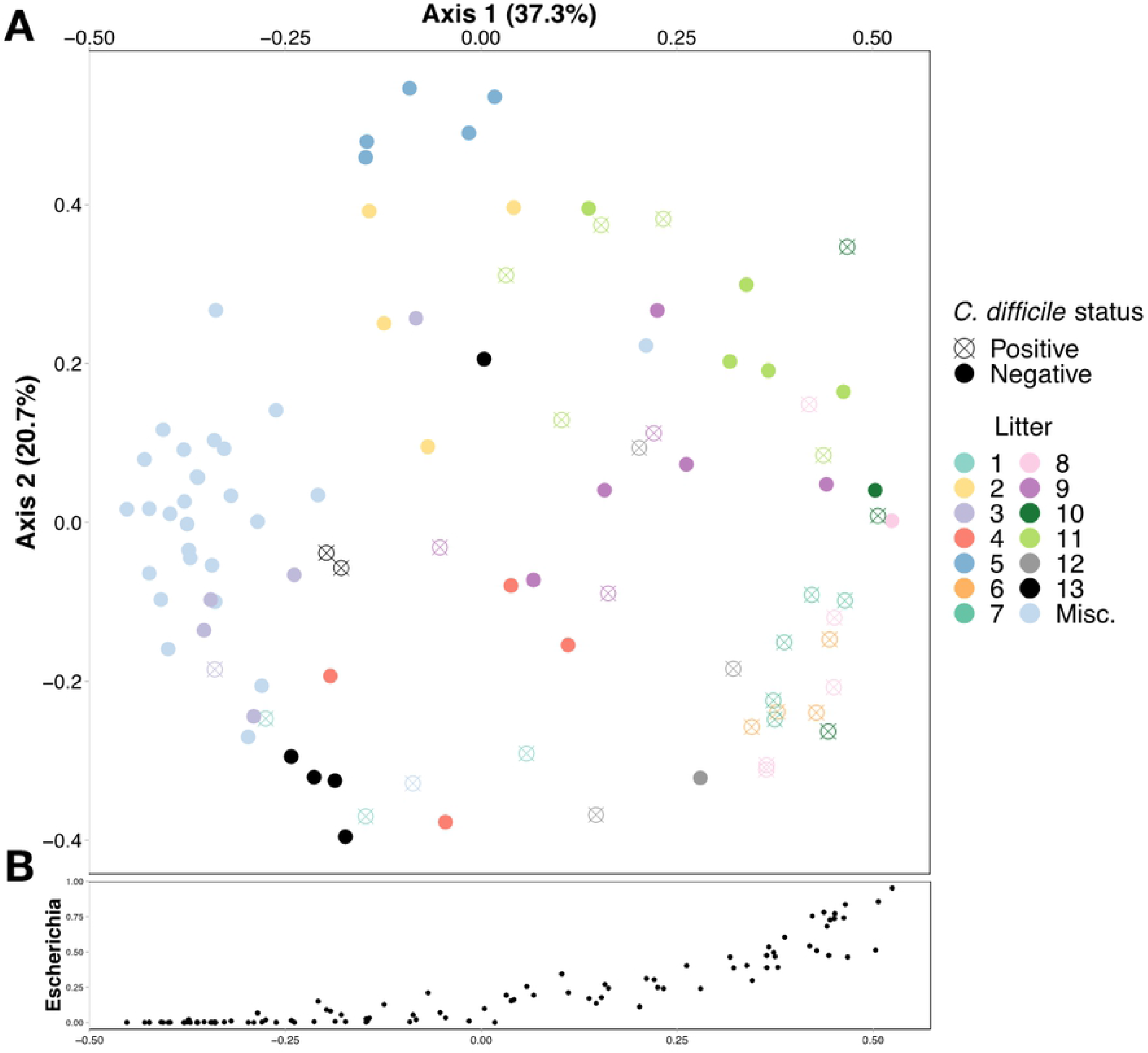
**A.** Bray-Curtis principal component analysis shows clustering of fecal samples from puppies in the greater Philadelphia area by *C. difficile* colonization status and by litter. Fecal samples labeled “Misc.” are from older weaned puppies that were no longer in litters and resided with their owner. **B.** Relative abundance of the genus *Escherichia* increases along the x-axis of the PCoA.

### Association between *C. difficile* status, age, litter, and microbiome diversity in unweaned puppies

To evaluate the effect of litter on the observed association between fecal bacterial community diversity and *C. difficile* colonization, we restricted analysis to the 70 unweaned puppies from 13 litters for which litter data were available. Seven of these litters consisted of a mix of colonized and non-colonized puppies, whereas in six of these litters, all of the puppies were of the same colonization status. When controlling for litter, *C. difficile* status had no effect on microbial alpha diversity (p=0.5468). Among these unweaned puppies, the litter explained most (67%, p= 1.0e-4) of the dissimilarity between bacterial communities, and colonization with *C. difficile* was no longer significantly correlated with microbiome composition (p > 0.1). PCoA analysis showed distinct clustering within most litters, but not necessarily by colonization status within a litter (Supplemental Figure 1).

**Supplemental Figure 1.** Principal component analysis plot showing clustering of fecal samples from seven litters of puppies in the greater Philadelphia area where puppies within litters had different *C. difficile* colonization status

## Discussion

Asymptomatic carriage of *C. difficile* is common in young animals of many species, including humans, dogs, pigs, and horses. In people, colonization with *C. difficile* has been shown to be associated with altered gut microbial diversity (17, 19, 36–38), but no studies have examined this association in dogs. We found that the association between lower bacterial community diversity and *C. difficile* colonization was statistically significant even when accounting for age, and certain bacterial taxa were preferentially associated with *C. difficile* colonization.

As has been found in other studies (14, 25), both colonization with *C. difficile* and reduced gut microbial diversity in puppies were significantly associated with young age. Similar associations have also been found in human studies (17, 19, 36, 39). However, within litters, this association was no longer significant. Puppies of a same litter are exposed to the same environment, consume the same diet (i.e., dam’s milk), and are cophrophagic. It is therefore not surprising that similar gut microbial communities are seen among puppies of a litter, as we found and as was found in a previous study of 30 German Shepherd litters (40). Microbial communities, presumably along with *C. difficile*, are likely shared among littermates. However, even within litters, we noted heterogeneity in the fecal microbiome (Figure 5) and in colonization status. In more than half of the litters (7/13), there were colonized and non-colonized puppies, suggesting that either our sample sizes were too small to detect a significant association between colonization status and microbial diversity, or other unmeasured factors were associated with colonization. The heterogeneity in the fecal microbiome within a litter may be analogous to the cage effect in mice studies (41, 42), where significant interindividual differences in intestinal microbiota were seen among mice within a cage, even though they were bred and raised in highly controlled similar conditions.

While the association between gut microbial diversity and *C. difficile* colonization status did not attain statistical significance within a litter, it is likely that features of the gut microbiome nevertheless contribute to the establishment and persistence of *C. difficile*. We found *C. difficile*-positive samples to be enriched with members of the *Escherichia, Bacteroides, Enterococcus* and *Parabacteroides* genera, and *C. difficile*-negative samples with members of the *Prevotella, Megamonas*, and *Streptococcus* genera. Almost identical trends were found for taxa of the *Escherichia, Parabacteroides, Enterococcus, Prevotella* and *Megamonas* genera in one study comparing *C. difficile* non-colonized, asymptomatically colonized and infected human adults (19), and for taxa of the *Parabacteroides, Prevotella, Paraprevotella* and *Enterococcus* genera in another study of non-colonized and colonized adults (38). Similar findings were found for the *Bacteroides* genera in a study of human infants *(39)*. In particular, increased relative abundances of taxa from the *Parabacteroides* and *Enterococcus* genera are thought to be the result of a blooming phenomenon associated with reduced ecological niche competition in people with CDI (38, 43, 44).

Among unweaned puppies, we found that noncolonized puppies had higher relative abundances of taxa from the *Clostridia* genera compared to colonized puppies. Consistent with this finding, other studies have postulated that bacterial species that are phylogenetically related to *C. difficile* and share niches and compete for similar resources could provide colonization resistance against toxigenic *C. difficile* (45, 46). In fact, colonization with non-toxigenic *C. difficile* has been shown to prevent infection with toxigenic *C. difficile* in hamsters and people following administration of antibiotics (46–48).

In contrast to our findings, one study showed that noncolonized human infants had lower relative abundance of taxa from the *Escherichia* genera than colonized infants (17), while several other studies found *Bacteroides* spp in greater relative abundance in non-colonized human infants, children and adults *(19, 36, 49, 50)*. It is unclear why these discrepancies were observed in our study. Both *Bacteroides* spp, which are used as markers of a healthy gut in people (50), and *E. coli* are found in the feces of healthy puppies (25, 51). *Bacteroides* spp are found in increasing relative abundance with increasing age, while *E. coli* levels are significantly higher in younger (less than 21 days) puppies than in older (greater than 42 days) puppies (25). Our findings underscore that puppies colonized with *C. difficile* nevertheless retain much of the gut microbiota of healthy animals and point to possible species-specific differences in the impact of *C. difficile* on the gut microbiome.

While some of the general trends were similar in our study and in several human studies, it is important to note that GI microbiota differ significantly by species, and extrapolation from human to animals is not always possible or prudent. In one study, for example, microbial groups associated with *C. difficile* colonization status were significantly different for people and poultry (52). However, the canine gut microbiome has been shown to be more similar to the human gut microbiome than that of pigs and mice (53, 54), perhaps due to their shared environments and diets, which might be why we observed similar microbiological trends in puppies and people.

The large proportion of puppies colonized with *C. difficile* has important implications for the potential zoonotic transmission of this organism. While it is likely that a puppy’s litter (and resultant environmental exposures) is the main determinant of colonization status, it is also likely that the puppy’s microbiome has an effect. The small number of puppies in each litter and the limited number of litters with colonized and non-colonized puppies precluded us from establishing whether the effect was statistically significant, but microbial community signatures that were consistent with what has been observed in people suggest that the microbiome has a role to play in colonization resistance. The protective role of the gut microbiome is particularly important when considering the fact that many puppies sold in pet stores (up to 95%) receive prophylactic antibiotics prior to shipping, as was recognized in a recent outbreak of Campylobacteriosis associated with puppies in pet stores (55). This could result in gastrointestinal dysbiosis and a resultant predisposition to harboring pathogens such as *C. difficile*. More research is needed to (1) better understand the interaction between the gut microbiome and colonization and infection with *C. difficile* in dogs, especially at the level of the litter; (2) define the relationship between dog-colonizing *C. difficile* strains and human colonizing strains; and (3) understand how interventions that reduce colonization in human pets may impact human disease prevention.

## Conclusions

We found that colonization with *C. difficile* is associated with reduced gut microbiome diversity in puppies, even when adjusting for the puppy’s age, and that there were differentially-abundant taxa in *C. difficile*-positive and *C. difficile*-negative fecal samples that may be permissive in promoting the colonization and establishment of *C. difficile*. Though this effect was not observed at the level of the litter, and even though the litter explained a large proportion of the gut microbiome diversity, heterogeneity in the gut microbiome and in *C. difficile* colonization within litters was observed in more than half of the litters, suggesting that the gut microbiome and potentially other unmeasured factors contribute to colonization resistance against *C. difficile* in puppies.

## Authors’ contributions

ASB: data curation, methodology, formal analysis, investigation, software and writing – original draft preparation and review and editing, visualization

DB: investigation, methodology

BJK: conceptualization, formal analysis, methodology, writing – review and editing

DJK: investigation, methodology, writing

DPB: Supervision, writing, resources

RNB: Supervision, resources

LER: conceptualization, data curation, funding acquisition, investigation, methodology, project administration, resources, supervision, writing – original draft preparation and review and editing

